# Ploidy variation modulates outbreeding response and promotes mating system evolution in a selfing plant lineage

**DOI:** 10.1101/2025.04.15.648901

**Authors:** Carlos Olmedo-Castellanos, Ana García-Muñoz, Camilo Ferrón, Mohammed Bakkali, A. Jesús Muñoz-Pajares, Mohamed Abdelaziz

## Abstract

- Outbreeding response, the phenotypic differences observed between selfed parental lines and their outcrossed offspring, can influence the evolution of selfing strategies. However, such effect remains poorly understood in non-crop species. We investigated the phenotypic outbreeding response variation across ploidy levels in *Erysimum incanum*, a predominantly selfing plant complex with diploid, tetraploid, and hexaploid populations distributed across the Iberian Peninsula and Morocco.
- We performed controlled within-population crosses to generate offspring with varying heterozygosity levels across ploidy types. We quantified individual, flower, and reproductive traits, and we estimated fitness components, and assessed trait modularity and phenotypic integration to see how heterozygosity affects trait coordination.
- Tetraploid showed the strongest and most consistently positive outbreeding responses, particularly in gamete production. Trait-specific outbreeding responses were positively associated with fitness across ploidy levels. Increasing heterozygosity was linked to a reduction in phenotypic integration, suggesting a loosening of trait correlations.
- Our results show that outbreeding response is ploidy-dependent and functionally connected to fitness. This suggests it may act as a selective force promoting outcrossing in highly inbred lineages. We suggest that outbreeding response is a dynamic and evolvable trait, with implications for mating system transitions and diversification in selfing plant populations.

## INTRODUCTION

Mating systems in flowering plants range from self-pollination (selfing) to cross-pollination (outcrossing), with these two extremes considered the most evolutionarily stable strategies (Laenen et al., 2018). However, in many predominantly selfing species, individuals can still receive pollen from unrelated conspecifics, allowing for occasional outcrossing. A shift in the mating system may occur if outcrossed offspring exhibit higher fitness than those produced by selfing—a studied scenario by simulations in contact zones between formerly isolated lineages (Harkness et al., 2019). Hence, following the terminology used by Whitlock et al. (2013), we define outbreeding response as the net phenotypic or fitness change observed in the offspring produced by crossing two genetically distinct selfing individuals. Since selfing individuals are typically highly inbred and exhibit low heterozygosity, any outbreeding response is largely driven by heterozygosity increases (Kumar et al., 2020). This response may lead to either heterosis or outbreeding depression, depending on whether the fitness effects are positive or negative (Oakley et al., 2015; Soto et al., 2023). Notably, both outcomes can occur across different traits or generations within the same lineage, as the strength and direction of the response depend on complex genetic, epigenetic, phenotypic, and environmental interactions (Govindaraju, 2019). Ultimately, the key evolutionary question centers on the net effect of outcrossing on fitness, and the adaptive relevance of outbreeding response remains incompletely understood (Frankham, 2015).

While outbreeding effects have been widely examined in agricultural systems (Labroo et al., 2021; Hochholdinger & Baldauf, 2018; Liu et al., 2022), their evolutionary implications in natural systems remain unclear. Despite being a topic of interest since Darwin (1876), the evolutionary role of outbreeding response remains elusive due to its dependence on genetic background, ecological context, and especially genetic distance between parents (Van Dooren, 2000). In general, outbreeding response tends to increase with parental genetic distance, though excessive divergence can result in hybrid breakdown due to incompatibilities (Rehman et al., 2021). Conversely, crosses between closely related yet inbred individuals may exhibit heterosis due to masked deleterious alleles (Rhode & Cruzan, 2005). This gives rise to the concept of an “optimal outcrossing distance”, where fitness benefits are maximized while both inbreeding and outbreeding depression are avoided (Edmands, 1999; Chung et al., 2023).

Importantly, not all traits respond to outbreeding in the same way. Traits not directly related to fitness often show stronger heterotic effects, while fitness-related traits tend to exhibit weaker or even negative responses, likely due to their complex epistatic interactions and reduced heritability (Whitlock et al., 2013). Moreover, differences in how fitness is measured—*i.e.*, whether via growth, reproduction, or survival—can influence interpretations of heterosis and outbreeding depression. From an evolutionary standpoint, it is essential to focus not merely on phenotypic changes but on their relationship with reproductive success and long-term adaptability too.

Polyploidy is widespread in plant lineages and adds another layer of complexity to outbreeding responses (Renny-Byfield et al., 2014). Polyploids tend to harbor higher levels of heterozygosity, exhibit stronger heterotic effects (Chen, 2010), and display altered gene expression and trait variance compared to diploids (Riddle et al., 2010; Bansal et al., 2012). These genomic characteristics influence phenotypic integration—the degree to which traits co-vary—and may contribute to wider phenotypic spaces and greater evolutionary potential (Lopez-Jurado et al., 2022). Additionally, heterozygosity increase and altered gene interactions following outcrossing can weaken trait covariation patterns, further reshaping phenotypic architecture (Palao et al., 2011). Theoretical models suggest that phenotypic integration is positively associated with inbreeding depression and negatively correlated with the potential for heterosis (Vasseur et al., 2019), implying that low integration may facilitate transitions in fitness landscapes.

The *Erysimum incanum* complex provides a compelling system to investigate how outbreeding responses are modulated by ploidy and trait architecture. This selfing species, native to the Iberian Peninsula and Morocco, exhibits diploid, tetraploid, and hexaploid cytotypes (Nieto Feliner, 1993). Its short monocarpic life cycle facilitates the measurement of trait variation and fitness across generations. Furthermore, polyploidy within *E. incanum* allows for the exploration of how genome duplication alters the adaptive consequences of occasional outcrossing in a predominantly selfing background.

If outbreeding response leads to higher fitness in cross-fertilized offspring, it may act as a catalyst for transitions from selfing to outcrossing. In turn, the genomic flexibility conferred by polyploidy may amplify this potential by enabling more diverse phenotypic outcomes. Furthermore, reduced phenotypic integration may facilitate movement through new adaptive landscapes. To test these ideas, we (1) quantify outbreeding response, (2) evaluate how ploidy and trait modules affect its magnitude, (3) investigate the relationship between phenotypic integration and outbreeding response, and (4) assess whether outbreeding response acts as a selection target.

To this end, we performed controlled crosses among genetically distinct selfing maternal lines of *E. incanum*, belonging to the same population and ploidy level, to generate outcrossed offspring with increased heterozygosity. These were compared with offspring produced via self-fertilization. Additionally, we established second-generation selfed lines from the outcrossed individuals to evaluate trait reversibility and persistence. Through this framework, we aim to determine whether outbreeding enhances fitness and may thus contribute to evolutionary shifts in mating systems.

## MATERIAL AND METHODS

### Study system

*Erysimum* L. is a highly diverse genus within the Brassicaceae family, distributed across Eurasia, North Africa, and parts of North and Central America (Al-Shehbaz et al., 2006). One of its species, *Erysimum incanum* (Fig. 1), is considered a species complex, comprising annual and monocarpic taxa and subspecies inhabiting the Western Mediterranean Basin (Nieto-Feliner & Clot, 1993; Abdelaziz et al., 2019). This complex includes three ploidy levels: diploids (2n = 2x = 16 chromosomes), tetraploids (2n = 4x = 32), and hexaploids (2n = 6x = 48) (Nieto-Feliner & Clot, 1993; Abdelaziz et al., in prep.). Diploids occur in a vicariant distribution across the Rif and Pyrenees Mountains (Fig. 1b), while tetraploids are found in the southwestern Iberian Peninsula and the Middle Atlas Mountains (Nieto-Feliner & Clot, 1993; Fennane & Ibn-Tattou et al., 1999). In contrast, hexaploids are restricted to the southernmost ranges of Morocco (High Atlas and Anti-Atlas; Abdelaziz et al., in prep.). The *E. incanum* complex is predominantly autogamous. Its flowers are small, hermaphroditic, self-compatible, and exhibit an anther-rubbing mechanism that promotes efficient self-pollination (Abdelaziz et al., 2019).

**Fig. 1.**
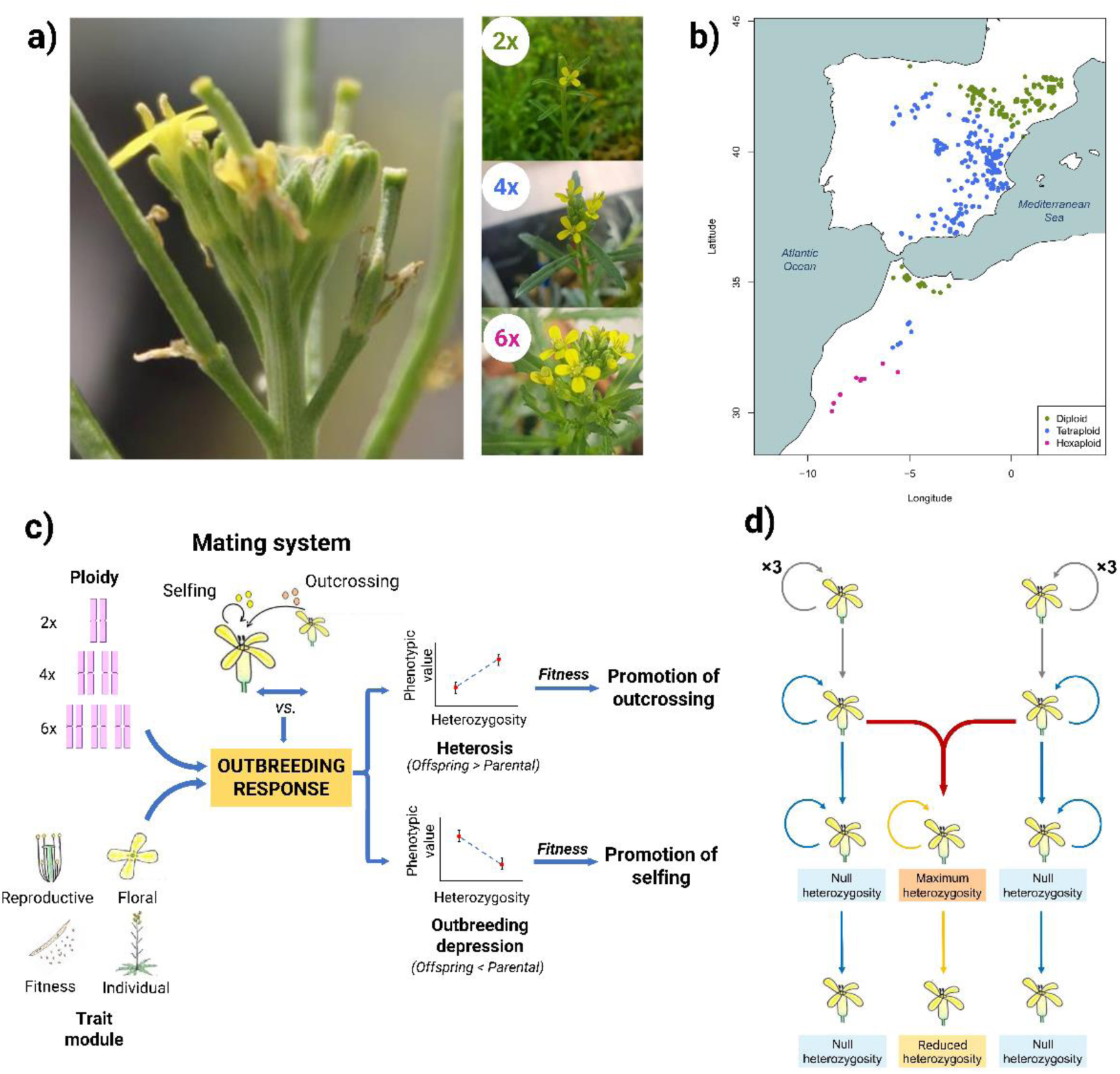
Experimental framework to study outbreeding response. **a.** *Erysimum incanum* flowers and representative individuals of each ploidy level. **b.** Geographic distribution of *E. incanum* populations across the Iberian Peninsula and Morocco. **c.** Conceptual workflow of the experimental design. Is outbreeding response a mechanism that promotes outcrossing? If a positive outbreeding response increases fitness, then outcrossing may be favoured over selfing. This response may be influenced by ploidy level and trait modularity. The effect of cross-fertilization between selfing parental lines could vary depending on ploidy and trait module, and the associated fitness changes may favour a shift toward either selfing (negative response) or outcrossing (positive response). **d.** Experimental design of greenhouse crosses to generate offspring with different heterozygosity levels. We established three experimental groups: (1) selfed offspring with null heterozygosity, derived from self-pollination; (2) outcrossed offspring with maximum heterozygosity, resulting from controlled cross-fertilizations between inbred lines; and (3) selfed offspring from the previous outcrossed group, displaying intermediate heterozygosity, produced via controlled self-pollination.

### Experimental design and phenotypic measurements

We sowed seeds from six wild populations (Table 1) in the greenhouse facilities in the Faculty of Sciences (University of Granada) for five consecutive generations to reduce residual heterozygosity resulting from potential previous outcrossing events. We then conducted controlled crosses among different maternal lines within each population to generate three heterozygosity groups: (i) null heterozygotes, derived from selfing; (ii) maximum heterozygotes, produced by controlled outcrossing between inbred lines; and (iii) reduced heterozygotes, obtained by selfing the maximum heterozygote offspring.

**Table 1.** Geographical coordinates of natural populations of *Erysimum incanum* used for experiment.

| Ploidy | Population | Location | Coordinates |
| --- | --- | --- | --- |
| Diploid | *Ei12* | Rif (Morocco) | 35° 11' 8" N<br>5° 13' 19" W |
| Diploid | *Ei13* | Pyrenees (Spain) | 41° 58' 20" N<br>0° 46' 43" W |
| Tetraploid | *Ei08* | Middle Atlas (Morocco) | 33° 27' 06" N<br>4° 57' 38" W |
| Tetraploid | *Ei09* | Guadix (Spain) | 37° 20' 18" N<br>3° 02' 38" W |
| Hexaploid | *Ei16* | Anti-Atlas (Morocco) | 30° 50' 04" N<br>8° 23' 38" W |
| Hexaploid | *Ei18* | Anti-Atlas (Morocco) | 30° 14' 16" N<br>8° 10' 37" W |

For each ploidy level, we first inter-crossed highly inbred maternal lines to generate maximum heterozygotes. We then allowed three generations of unassisted selfing to produce null heterozygotes. Finally, we performed controlled selfing on maximum heterozygotes to obtain reduced heterozygotes (Fig. 1d). In total, we obtained 405 diploid plants (257 null, 53 maximum, and 95 reduced heterozygotes), 764 tetraploid plants (457 null, 69 maximum, and 238 reduced heterozygotes), and 453 hexaploid plants (252 null, 112 maximum, and 89 reduced heterozygotes).

We measured flower traits during anthesis using one flower per plant and a digital calliper (±0.1 mm accuracy; Dexter, Alcobendas, Madrid). Measured traits included petal length (maximum length from the edge to the curving point), corolla diameter (distance between opposite petals), and corolla tube length (distance from sepal base to the corolla opening). We also measured the lengths of long and short filaments and the height of the style (from its base to the stigma), allowing us to calculate herkogamy (i.e., the vertical distance between long stamens and the style).

To assess male reproductive investment, we collected half of the stamens per flower and preserved them in 100 µL of 70% ethanol. We estimated pollen quantity per flower using a Multisizer Coulter Counter 3™ (Beckman Coulter, Pasadena, USA), provided by the morphometric laboratory at the Centro de Investigación, Tecnología e Innovación (CITIUS), University of Sevilla (Spain).

We collected mature fruits and counted viable seeds, aborted seeds, and unfertilized ovules per fruit to estimate female reproductive investment. We also evaluated seed germination rate by sowing 10 seeds per pot (per heterozygosity group) and calculating the proportion that germinated. Additional measured variables included the number of leaves and fruits per plant, the number of seeds per fruit (average from four fruits), the height of the main flowering stalk, the number and diameter of all stalks, and the total number of flowers per plant.

### Statistical analysis

#### Outbreeding response

We grouped traits into four modules: (i) individual traits (height, number of stalks, stalk diameter, and number of flowers), (ii) flower traits (petal length, flower diameter, corolla length, long filament, short filament, and herkogamy), (iii) reproductive traits (mean number of pollen grains and ovules per flower), and (iv) fitness components (total pollen and ovule output per plant, fruit set, seed set, germination, survival, and number of leaves).

We calculated outbreeding response (OR) using a modified version of the inbreeding depression index (Johnston & Schoen, 1994):

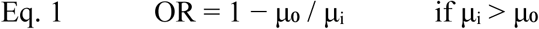

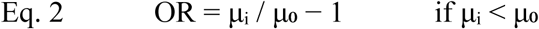

Where μ₀ represents trait values in null heterozygotes and μᵢ those in heterozygous individuals (either maximum or reduced). This index, ranging from −1 (maximum outbreeding depression) to +1 (maximum heterosis), allows standardized comparisons across traits and ploidy levels.

To assess the statistical significance of outbreeding responses, we used bootstrap resampling (n = 100,000) to calculate 95% confidence intervals. We also computed trait correlation–covariance matrices for each ploidy level to evaluate inter-trait relationships. To test whether outbreeding response varied by ploidy level or trait module, we applied the Scheirer–Ray–Hare test, a non-parametric alternative to factorial ANOVA.

#### Linear regression of individual contribution to outbreeding response on fitness

To evaluate whether outbreeding responses are subject to natural selection, we calculated each individual plant’s contribution to outbreeding response by comparing its trait values (under maximum or reduced heterozygosity) with the population mean under null heterozygosity. We used the same formulae (Eq. 1 and Eq. 2) and then normalized the output using:

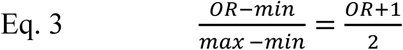

This transformation scales values from (−1,1) to (0,1), which allows interpreting contributions in a positive range (see Table S1 for whole detail).

We calculated fitness as the absolute product of fruit production and seed production, and we normalized it by the higher value in every ploidy level. For each trait, we performed linear regressions between the individual contribution to outbreeding response and fitness.

#### Phenotypic integration

We assessed phenotypic integration by quantifying the number of significant pairwise correlations among traits. We grouped individuals by ploidy (diploid, tetraploid, hexaploid) and heterozygosity level (null, maximum, reduced). For each group, we calculated the number of significant correlations involving each trait and derived the average number of correlations per trait–group combination.

## RESULTS

We evaluated the effect of cross-fertilization on phenotypic traits in *E. incanum* lines differing in ploidy (diploid, tetraploid, and hexaploid) and heterozygosity level (maximum, null, and reduced). Table 2 shows the outbreeding response across trait modules and heterozygosity levels, so that:

### Outbreeding response for maximum heterozygosity

Tetraploids exhibited the highest number of traits with significant outbreeding responses (14 out of 20), of which 13 were positive. In contrast, diploids and hexaploids each showed only 4 traits with significant responses: all 4 were positive in hexaploids, while 3 of 4 were negative in diploids (Table 2a).

**Table 2a.** Outbreeding response values for phenotypic traits in *Erysimum incanum* for maximum heterozygosity group. Significant values are highlighted in red (negative) and blue (positive). Bootstrapped confidence intervals were calculated based on 100,000 replicates.

| Module | Diploid | 95 % CI | Tetraploid | 95 % CI | Hexaploid | 95 % CI |
| --- | --- | --- | --- | --- | --- | --- |
| *Fitness* |  |  |  |  |  |  |
| Ovules/Plant | -0.041 | (-0.285, 0.239) | 0.403 | (0.163, 0.562) | 0.230 | (-0.042, 0.425) |
| Pollen/Plant | 0.156 | (-1.325, 0.566) | 0.687 | (0.528, 0.786) | 0.064 | (-0.290, 0.318) |
| Fruitset | 0.103 | (-0.014, 0.189) | 0.227 | (0.181, 0.269) | 0.039 | (-0.015, 0.088) |
| Seedset | 0.059 | (-0.015, 0.112) | 0.027 | (-0.061, -0.068) | -0.068 | (-0.174, 0.032) |
| Survival | 0.292 | (0.227, 0.347) | 0.015 | (-0.119, 0.127) | -0.084 | (-0.169, 0.009) |
| Germination | -0.473 | (-0.559, -0.380) | -0.210 | (-0.298, -0.120) | -0.181 | (-0.322, 0.002) |
| Number of leaves | -0.223 | (-0.326, -0.114) | -0.045 | (-0.132, 0.039) | 0.024 | (-0.172, 0.199) |
| *Reproductive* |  |  |  |  |  |  |
| Ovules/Flower | -0.019 | (-0.173, 0.147) | 0.371 | (0.291, 0.437) | 0.162 | (0.076, 0.238) |
| Pollen/Flower | 0.179 | (-0.227, 0.421) | 0.554 | (0.445, 0.638) | 0.192 | (0.042, 0.316) |
| *Individual* |  |  |  |  |  |  |
| Plant height | 0.003 | (-0.184, 0.151) | 0.473 | (0.410, 0.530) | 0.115 | (-0.013, 0.224) |
| Number of stalks | -0.187 | (-0.438, 0.121) | -0.138 | (-0.326, 0.073) | -0.148 | (-0.328, 0.078) |
| Stalk diameter | 0.107 | (-0.106, 0.271) | 0.396 | (0.304, 0.474) | -0.021 | (-0.148, 0.121) |
| Number of flowers | -0.147 | (-0.363, 0.099) | 0.196 | (-0.006, 0.348) | -0.063 | (-0.259, 0.175) |
| *Flower* |  |  |  |  |  |  |
| Petal length | -0.089 | (-0.163, -0.003) | 0.135 | (0.089, 0.178) | 0.015 | (-0.044, 0.068) |
| Flower diameter | -0.060 | (-0.122, 0.006) | 0.141 | (0.098, 0.182) | -0.030 | (-0.072, 0.014) |
| Corolla tube length | -0.017 | (-0.083, 0.062) | 0.130 | (0.068, 0.192) | 0.045 | (-0.028, 0.112) |
| Long filament length | 0.044 | (-0.018, 0.101) | 0.199 | (0.172, 0.225) | 0.042 | (0.008, 0.074) |
| Short filament length | -0.035 | (-0.115, 0.050) | 0.154 | (0.115, 0.190) | 0.043 | (-0.008, 0.090) |
| Style length | -0.169 | (-0.424, 0.081) | 0.210 | (0.175, 0.244) | 0.071 | (0.026, 0.115) |
| Herkogamy | -0.235 | (-0.480, 0.012) | -0.044 | (-0.087, 0.000) | 0.040 | (-0.016, 0.091) |

The significant responses observed in diploids and hexaploids were not always shared. All four positive traits in hexaploids were also significantly positive in tetraploids, with larger effect sizes in tetraploids. Among diploids, only germination showed a significant negative response also shared by tetraploids; two traits (survival and number of leaves) were significant only in diploids—survival positive and leaves negative—while petal length showed opposing significant responses in diploids (negative) and tetraploids (positive).

Considering trait modules, no fitness component showed significant response in hexaploids, while the mean number of ovules per plant, the mean number of pollen per plant and the seed set were positively significant only in tetraploids. Survival was significant (positive) only in diploids. Germination was significantly negative both in diploids and tetraploids, the number of leaves per plant was significantly negative only in diploids, and the seed set being the only fitness component trait not showing significant outbreeding response at any ploidy level.

In the reproductive trait module, both pollen and ovule production per flower showed significant positive response in tetraploids and hexaploids but not significant in diploids.

For individual traits, only stalk diameter and plant height showed significant positive response and only in tetraploids. The number of stalks and number of flowers showed no significant response at any ploidy level.

In the flower trait module, long filament and style length had congruent positive responses in tetraploids and hexaploids. Flower diameter, corolla tube length, and short filament length showed positive responses exclusively in tetraploids. Petal length had opposing responses in diploids and tetraploids. Herkogamy showed no significant outbreeding response in any ploidy level.

Notably, no trait exhibited a significant response across all three ploidy levels. Traits such as seed set, number of stalks, number of flowers, and herkogamy did not show any significant response in any group.

### Outbreeding response for reduced heterozygosity

Diploids exhibited the highest number of traits with significant negative outbreeding responses (14 of 16 significant traits; Table 2b). Tetraploids and hexaploids each had 12 traits showing significant response, with 7 being negative in both—but affecting different traits in each. Most traits that show significantly negative response in diploids were either also negative or non-significant in the other two ploidy levels. However, flower diameter showed a significantly positive response in both tetraploids and hexaploids, despite being close to zero.

**Table 2b.**
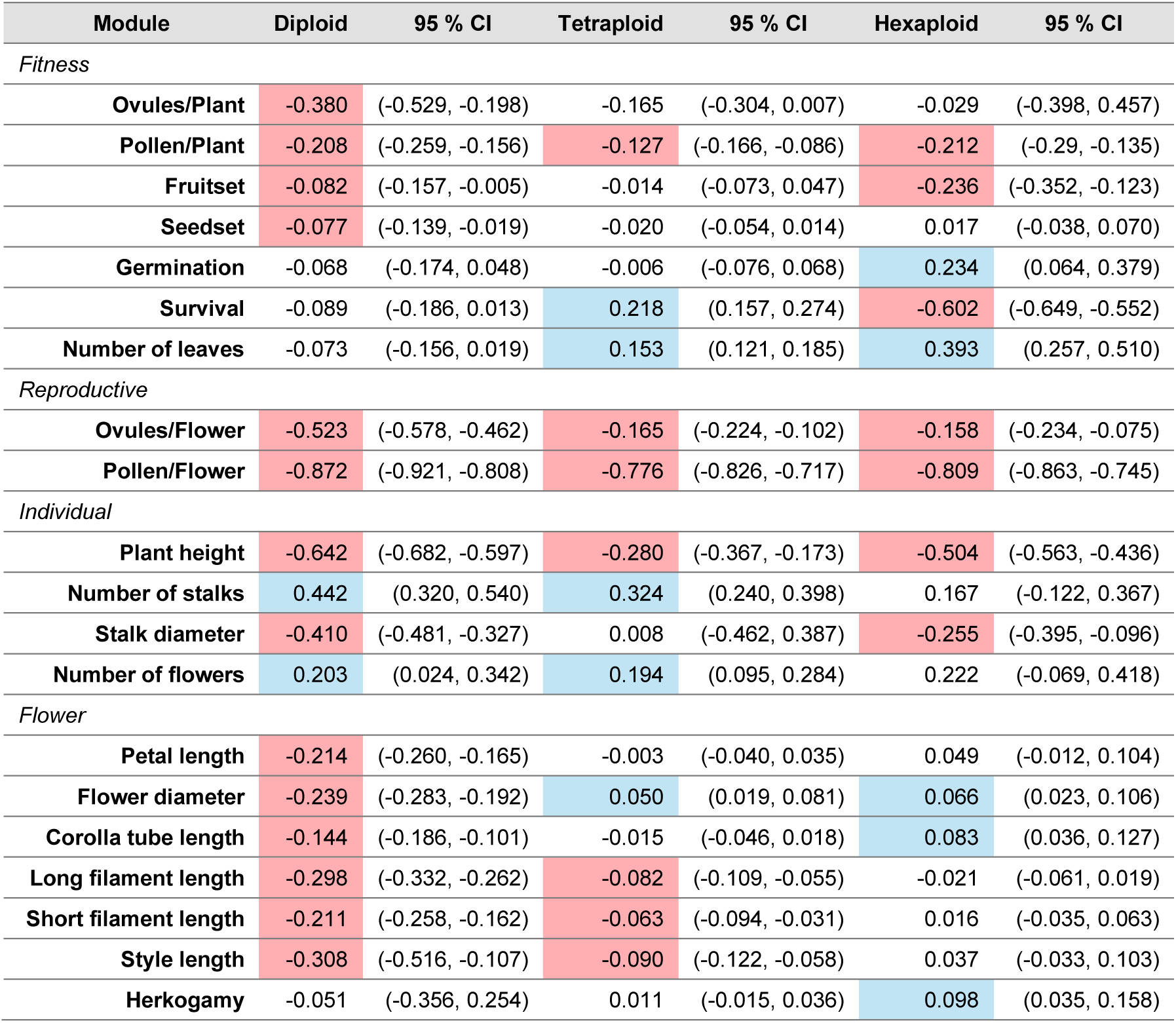
Outbreeding response values for phenotypic traits in *Erysimum incanum* for reduced heterozygosity group. Significant values are highlighted in red (negative) and blue (positive). Both heterozygosity levels are compared against the null heterozygosity group (selfed offspring). Bootstrapped confidence intervals were calculated based on 100,000 replicates.

For individual traits, stalk diameter and number of flowers showed significantly positive responses in diploids and tetraploids only. Reproductive traits showed consistently significant negative responses across all ploidy levels. In contrast, flower traits showed positive significant responses in hexaploids for three traits. Regarding fitness-related traits, ovule and pollen production per plant, fruit set, and seed set all had significantly negative responses. In contrast, traits linked to survival showed positively significant response in one or both polyploid levels (e.g., germination in hexaploids, survival in tetraploids, and number of leaves in both).

### Effects of ploidy and trait module on outbreeding

A Scheirer–Ray–Hare test (Fig. 2) revealed that at maximum heterozygosity, only ploidy had a significant effect on outbreeding response (H = 12.41; df = 2; p < 0.05). While, at reduced heterozygosity, both ploidy (H = 7.53; df = 2; p < 0.05) and trait module (H = 10.27; df = 3; p < 0.05) significantly affected outbreeding response.

**Fig. 2.**
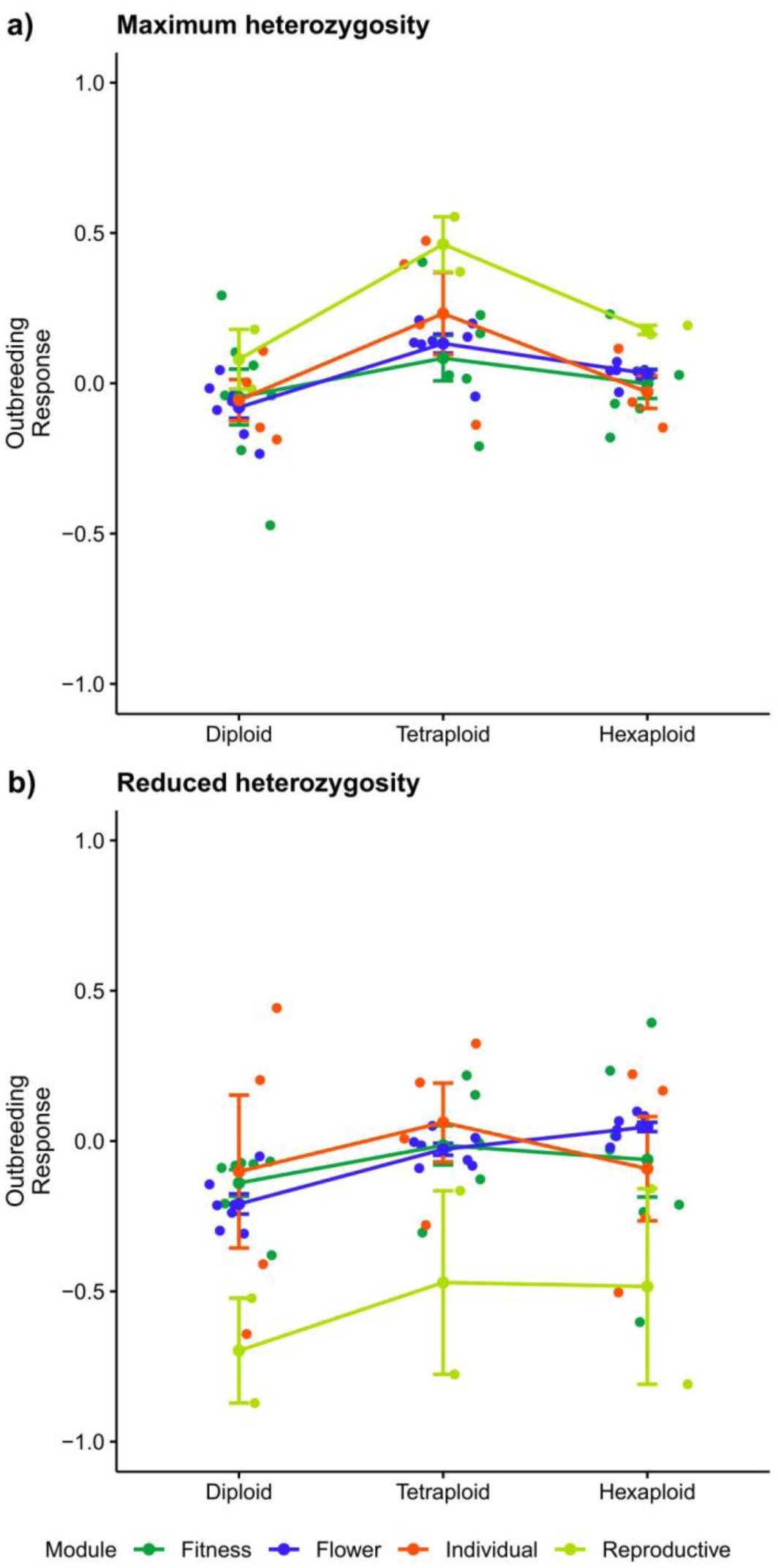
Variation in outbreeding response across trait modules and ploidy levels in *Erysimum incanum* under maximum and reduced heterozygosity. Each colour corresponds to a different trait module. Dots represent outbreeding response values for individual traits, while central dots indicate the mean response for each trait module. Significance was assessed using the Scheirer–Ray–Hare test. **a.** Outbreeding responses under maximum heterozygosity. **b.** Outbreeding responses under reduced heterozygosity.

### Fitness and outbreeding response

We estimated the individual contribution to outbreeding response for each trait, heterozygosity, and ploidy level to evaluate its relationship with fitness (Table 3). We found that all significant relationships were positive, except for survival in diploids with reduced heterozygosity, seed germination in diploids and hexaploids with reduced heterozygosity and herkogamy for tetraploids with reduced heterozygosity. The traits with the most significant relationships were the total amount of ovules, the number of flowers, and the total amount of pollen per plant. Our results show that a higher proportion of individual contribution to outbreeding response in these traits means a higher individual fitness.

**Table 3.** Linear regression coefficients of individual contributions to outbreeding response for each trait in *Erysimum incanum*, in relation to individual fitness (calculated as the product of seed set and fruit set). Regressions were performed separately for each ploidy level and heterozygosity comparison (maximum or reduced vs. null). Significance levels: **** *p* < 0.0001; *** *p* < 0.001; ** *p* < 0.01; * *p* < 0.05.

| Ploidy | Diploid |  | Tetraploid |  | Hexaploid |  |
| --- | --- | --- | --- | --- | --- | --- |
| Heterozygosity | Maximum | Reduced | Maximum | Reduced | Maximum | Reduced |
| Ovules/Plant | 0.156**** | 0.119**** | 0.262**** | 0.086**** | 0.203**** | 0.203**** |
| Pollen/Plant | 0.141 | 0.054 | 0.304** | 0.033 | 0.342**** | 0.339* |
| Fruit set | 0.087* | 0.076** | 0.192 | 0.071**** | 0.095 | 0.128* |
| Seed set | 0.115 | 0.082* | 0.15 | 0.067**** | 0.128** | 0.181 |
| Survival | 0.065 | -0.046** | -0.041 | 0.03** | -0.002 | -0.121 |
| Germination | 0.034 | -0.047*** | 0.028 | 0.016 | 0.056* | -0.047* |
| Number of leaves | 0.062* | -0.029 | 0.016 | 0.015 | 0.046 | -0.039* |
| Ovules/Flower | 0.064 | 0.172**** | 0.193 | 0.094**** | 0.271**** | 0.166** |
| Pollen/Flower | -0.08 | -0.024 | -0.175* | -0.025** | 0.156 | 0.018 |
| Plant height | 0.143**** | 0.189**** | 0.307* | 0.084**** | 0.115*** | 0.316* |
| Number of stalks | 0.126**** | 0.039** | 0.147**** | 0.007 | 0.149**** | 0.128** |
| Stalk diameter | 0.148**** | 0.025 | 0.204** | 0.065**** | 0.163**** | 0.111 |
| Number of flowers | 0.138**** | 0.06**** | 0.191**** | 0.071**** | 0.155**** | 0.141*** |
| Petal length | 0.014 | 0.021 | -0.04 | 0.039* | 0.117** | 0.009 |
| Corolla tube diameter | 0.022 | 0.055* | -0.121 | 0.057*** | 0.159** | 0.014 |
| Corolla length | 0.11 | 0.023 | -0.106 | 0.032 | 0.138*** | 0.041 |
| Long filament length | 0.186** | 0.016 | 0.323 | 0.042* | 0.245*** | 0.09 |
| Short filament length | 0.116* | -0.035 | 0.161 | 0.043** | 0.135** | 0.072 |
| Style length | 0.15* | 0.041 | 0.163 | 0.022 | 0.199** | 0.104* |
| Herkogamy | -0.01 | 0.027 | -0.057 | -0.034 | -0.04 | 0.134* |

### Phenotypic integration

We estimated trait integration and evaluated the effects of heterozygosity and ploidy levels (Fig. 3A, see Fig. S1 for separated traits). Overall, outcrossed offspring showed less integrated phenotypes compared to selfed individuals (Fig. 3B).

**Fig. 3.**
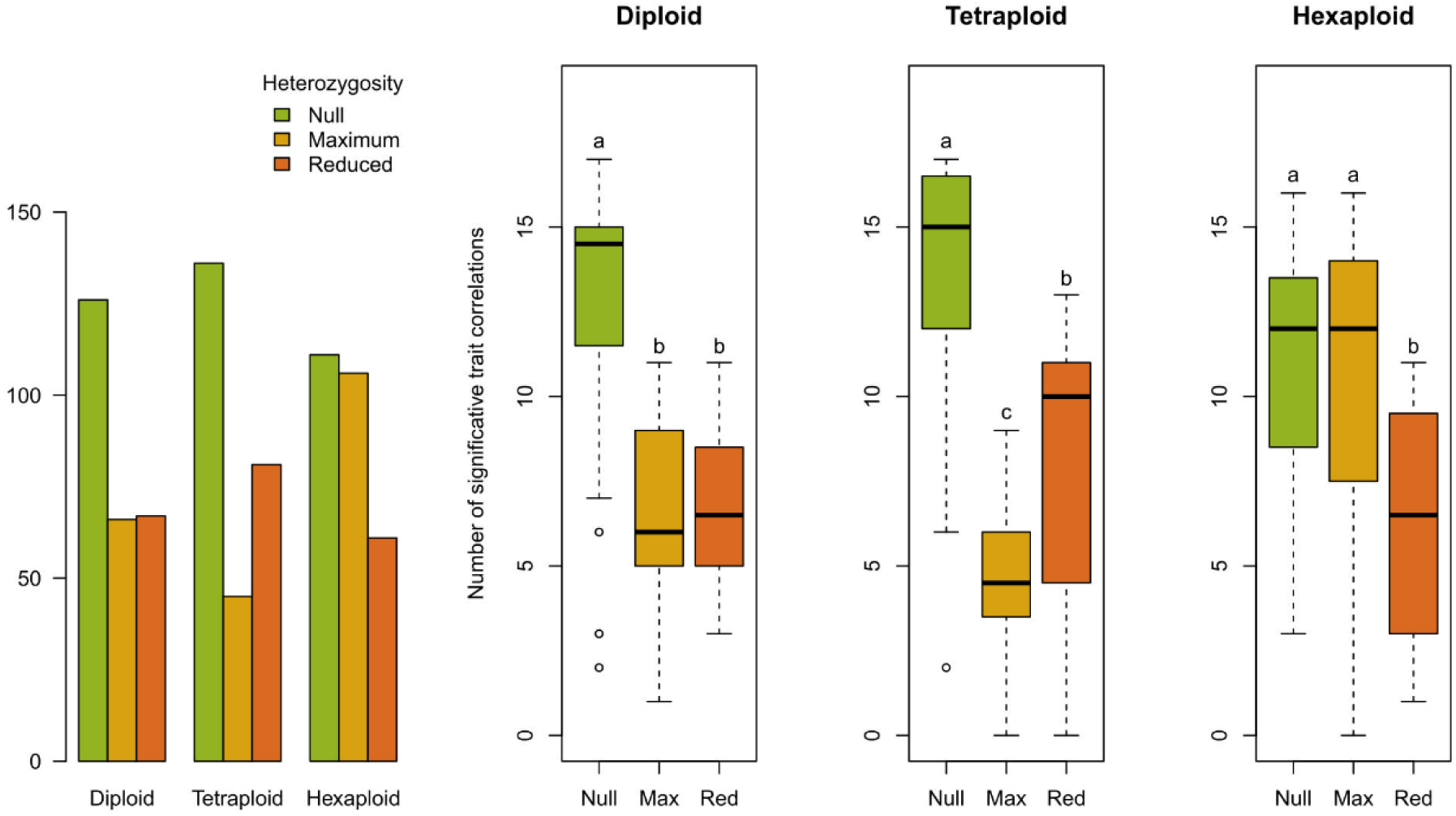
Patterns of phenotypic integration *in Erysimum incanum*. **a.** Total number of significant pairwise correlations (i.e., integrations) among traits. **b.** Mean number of significant correlations per phenotypic trait. Data are grouped by ploidy level and heterozygosity class. Significance was tested for differences between heterozygosity levels within each ploidy group using t-tests.

We observed a generally negative relationship between trait integration and heterozygosity level. Null-heterozygosity plants had the highest number of significant trait correlations across all ploidy levels. At higher heterozygosity, diploids and tetraploids showed reduced integration; but in hexaploids, integration remained similar to null heterozygotes

Selfed tetraploids exhibited the highest number of significant correlations overall. Interestingly, while increased heterozygosity reduced integration in diploids and tetraploids, hexaploids maintained stable integration levels, suggesting a potential buffering effect of higher ploidy.

## DISCUSSION

Outbreeding response has been extensively studied in the context of agriculture and breeding (Paril et al., 2024; Schnabel & Springel, 2013). However, it may also play a key role in evolution, potentially acting as a driver of outcrossing when it results in enhanced offspring fitness. *Erysimum incanum*, a selfing species within Brassicaceae, is highly inbred and comprises populations with varying ploidy levels—diploid, tetraploid, and hexaploid.

To evaluate the impact of outcrossing on fitness, we performed controlled crosses between distinct selfing lines. We found a significantly positive outbreeding response (i.e. heterosis) in tetraploids after a single outcrossing event. This was followed by a decrease in heterozygosity in subsequent selfed offspring, which was often associated with outbreeding depression. Overall, trait modules and ploidy levels explained a greater proportion of variation than heterozygosity level per se. Notably, we also detected a direct positive relationship between outbreeding response and fitness. In parallel, heterozygosity was inversely correlated with phenotypic integration across all ploidy levels. These findings suggest that outbreeding response may play a role in promoting transitions between mating systems.

A meta-analysis by Whitlock et al. (2013) found that outbreeding tends to negatively affect fitness-related traits. However, other studies, especially in polyploids, have shown fitness benefits following outcrossing between genetically distinct populations (Rodger et al., 2024). In our case, within-population outcrosses between selfing lines also enhanced fitness components. Specifically, tetraploid outcrossed individuals produced a higher total number of pollen grains and ovules and exhibited greater reproductive allocation per flower than their selfed counterparts.

Although fitness components are diverse and often context-dependent (Vasseur et al., 2019), our findings contrast with predictions that polyploidization promotes selfing and asexuality. Instead, they support growing evidence that polyploidy can expand phenotypic potential (Clo, 2022), enhancing heterotic effects (Chen, 2010). Still, increased phenotypic variation may not always translate into fitness benefits (Hedrick et al., 2012). For instance, while traits such as plant size or architecture may benefit from greater plasticity (Fort et al., 2014), hexaploids—with even broader phenotypic space— may face greater difficulty achieving optimal fitness peaks. The underlying mechanisms are complex (Mackay et al., 2021), but phenotypic expression remains a valuable proxy for exploring the functional consequences of outcrossing and genome duplication (Lippman & Zamir, 2007).

In diploids, outbreeding response at maximum heterozygosity was not statistically significant, although it tended to be negative across most traits. This contrasts with the predominantly positive responses observed in tetraploids and hexaploids. The inverse trend in diploids may reflect fundamental differences in the genetic basis of heterosis and outbreeding depression. Specifically, interactions among alleles and regulatory networks may be more susceptible to disruption in diploids than in polyploids. Increased heterozygosity can destabilize well-established genetic architectures in inbred backgrounds (Fridman, 2015). Furthermore, long-term divergence among selfing diploid lineages may lead to the accumulation of cryptic genetic incompatibilities, resembling those observed in between-population crosses (Clo et al., 2021). This effect is expected to be less pronounced in polyploids, where additional chromosome sets may buffer against allelic incompatibilities.

The subsequent reduction in heterozygosity through selfing of previously outcrossed diploids resulted in outbreeding depression, reinforcing a stabilization of the selfing strategy and making reversal toward outcrossing less likely. In contrast, the positive outbreeding response in polyploids suggests that purging of deleterious alleles has not been fully effective, allowing beneficial heterozygous combinations to persist. This incomplete purging may be particularly relevant for traits contributing to fitness, such as flower productivity and gamete output.

Ploidy emerged as the most influential factor explaining variation in outbreeding response. While polyploidy initially narrows the scope of heritable variation, this effect tends to be transient (Clo, 2022). The trait module was also a significant predictor, particularly at the reduced heterozygosity level. Trait modules not only reflect functional groupings but also shared developmental pathways and gene networks (Armbruster et al., 2013), making them more likely to exhibit coordinated responses to genetic perturbations.

For example, flower traits often exhibited more consistent and directional outbreeding responses than ind traits. This is likely due to their central role in reproductive success, which subjects them to stronger selective constraints and lower additive genetic variance (Whitlock et al., 2013). In contrast, individual traits are typically more plastic and influenced by abiotic conditions (Opedal et al., 2023). These differences in genetic architecture and selective regimes may explain why some modules are more responsive to outbreeding than others.

Fitness was positively associated with outbreeding response for multiple fitness-related traits, including total ovule and pollen production and the number of leaves (Rhode & Cruzan, 2005). This relationship was also evident at the reduced heterozygosity level (Table 2), particularly in diploids, where decreased fitness accompanied reduced heterozygosity, suggesting that outcrossing benefits may persist even after a subsequent selfing event. Individuals exhibiting greater positive outbreeding effects had higher reproductive success, which could favour the spread of outcrossing genotypes. Because the relationship between outbreeding and reproductive performance is positive, outbreeding depression following the loss of heterozygosity may result in reduced fitness, reinforcing the advantage of maintaining or restoring heterozygosity through outcrossing.

Although outbreeding response and heterozygosity are conceptually distinct, both can influence fitness (David, 1998). While outbreeding response refers to the net change in trait values following outcrossing, its evolutionary relevance depends on its fitness consequences. In our study, the consistent positive effect of outbreeding response on fitness highlights its potential evolutionary importance. Moreover, even short-term increases in fitness could benefit populations during colonization or environmental transitions (Barker et al., 2018). To our knowledge, this is the first study to explicitly explore the link between outbreeding response and multiple fitness components across ploidy levels and trait categories.

Phenotypic integration decreased with increasing heterozygosity in diploids and tetraploids, but not in hexaploids. Trait integration is shaped by underlying genetic correlations, which are disrupted by heterozygosity. The relative magnitude of heterozygosity increases after decrease of outcrossing with the increase of ploidy level —being greater in diploids than in tetraploids, and lowest in hexaploids. Even so, all heterozygosity levels showed reduced integration relative to null heterozygosity groups. In highly inbred populations, recombination is limited, and allelic combinations are preserved over generations. This stability promotes higher trait integration, particularly in selfing species, where flower integration is a hallmark of selfing syndromes (Fornoni et al., 2016). Disruption of long-standing genetic linkages through heterozygosity may reduce trait integration, potentially allowing novel phenotypic combinations. Such effects have been observed in *Arabidopsis*, where admixture between divergent genotypes produced transgressive trait variation (Palacio-López & Molofsky, 2021). Interestingly, despite their broader combinatorial potential, hexaploids maintained relatively high trait integration, possibly reflecting an ancient origin that allowed time for the reestablishment of stable networks. This functional integration in hexaploids may act as a constraint on phenotypic space compared to tetraploids.

According to simulation studies (Harkness et al., 2019), heterosis alone may be insufficient to drive mating system transitions, as its effects tend to be transient. These models suggest that long-term shifts from selfing to outcrossing require additional evolutionary forces. Still, heterosis may act in synergy with other selective processes, contributing to transitions under the right ecological and genetic conditions (Draghi & Whitlock, 2015). In our study, outcrossing occurred between selfing lines within populations—likely involving limited genetic divergence—so the magnitude of heterosis was expected to be modest. Nevertheless, the observed effects highlight the fine-scale influence of outbreeding response, even among closely related genotypes.

Our results provide empirical evidence that ploidy variation modulates outbreeding response, with consequences for fitness and evolutionary potential. Differences in reproductive outcomes among cytotypes suggest that both heterosis and outbreeding depression are not fixed properties but are shaped by genomic context, including gene expression patterns, epigenetic regulation, and developmental buffering (Comai, 2005; Chen, 2007). These findings underscore how polyploidy can reshape genetic architecture, modulating the trade-off between genetic diversity and reproductive efficiency, and ultimately influencing plant adaptation and diversification (Van de Peer et al., 2017; Soltis et al., 2014).

Overall, our study demonstrates a stronger outbreeding response in tetraploids of *Erysimum incanum*. The positive relationship between outbreeding response and fitness, combined with lower phenotypic integration, suggests that tetraploids occupy a broader phenotypic space than diploids. This flexibility may enhance their capacity to adapt to new or fluctuating environments. While selfing offers short-term reproductive assurance, it may limit long-term evolutionary potential. Outcrossing becomes adaptive when it results in higher-fitness offspring, thereby favouring a shift in mating system. However, for such a shift to have evolutionary impact, it must be stabilized within populations over time. This work provides a first step toward understanding the evolutionary dynamics of mating system transitions in *Erysimum*. Future research will aim to explore additional factors that could promote outcrossing and determine whether such transitions can be maintained in natural populations.

## DECLARATIONS

### ETHICS APPROVAL AND CONSENT TO PARTICIPATE

Not applicable

### CONSENT FOR PUBLICATION

Not applicable

### AVAILABILITY OF DATA AND MATERIALS

The dataset generated and user during the current study are available from the corresponding author on reasonable request.

## ACKNOWLEDGMENTS

We thank Melissa Viveiros-Moniz, Yannire Vázquez Benítez, Esther Funes, Mireia Bustos and Modesto Berbel for lab assistance and paper discussion at the research network *BioChange* (biochangenet.org). We thank Xavier Picó from Estación Biológica de Doñana (CSIC, Spain) and Josselin Clo from Evo-Evo-Paleo Laboratory (CNRS/University of Lille, France) for his comments about methodology and writing. We also thank Luis Matías and María Jesús Ariza from the University of Sevilla for their helpful assistance and management to work in CITIUS labs (University of Sevilla).

## AUTHOR CONTRIBUTIONS

COC wrote the first draft of the manuscript, did the statistical analysis and designed figures and tables. AGM, CF, MB, AJMP and MA designed the experiments. AGM and CF developed greenhouse experimental work. AGM, CF and MA collected phenotypic data. MA, AJMP and MB got the funding and supervised all the work and contributed to the writing.

## Supporting Information

**Fig. S1.**
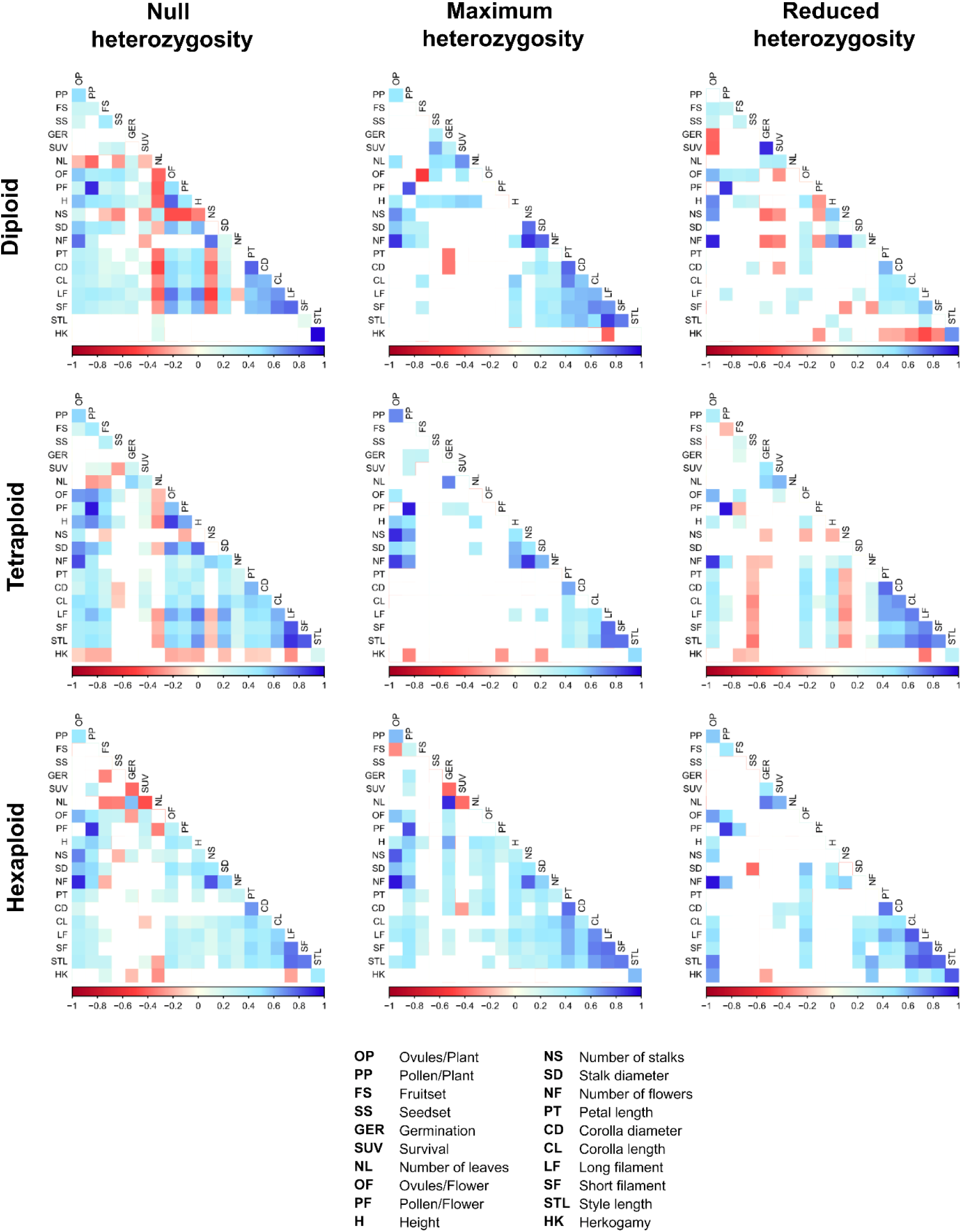
Significance levels of the correlations in *Erysimum incanum.* between traits for each ploidy (horizontal) and heterozygosity level (null, maximum and reduced; vertical). Red colour shows a negative correlation coefficients while blue shows positive correlation coefficients. Blank means not significant correlation.

**Table S1.** Mathematical development of the individual contribution to outbreeding response.

| Group level | Individual level |
| --- | --- |
| $1 - \frac{\mu_0}{\mu_i} \quad \text{if } \mu_i > \mu_0$ $\frac{\mu_i}{\mu_0} - 1 \quad \text{if } \mu_i < \mu_0$ | $1 - \frac{\mu_0}{i} \quad \text{if } i > \mu_0$ $\frac{i}{\mu_0} - 1 \quad \text{if } i < \mu_0$ |
| Result: Outbreeding response for a <u>group</u> in a given trait. | Result: Outbreeding response for an <u>individual</u> in a given trait. |
| *Linear normalization [0,1]* |  |
| **Individual contribution to outbreeding response** |  |
| $1 - \frac{OR'}{OR_i'} \quad \text{if } OR_i' > OR'$ $\frac{OR_i'}{OR'} - 1 \quad \text{if } OR_i' < OR'$ | |

